# Omicron BA.1 and BA.2 are antigenically distinct SARS-CoV-2 variants

**DOI:** 10.1101/2022.02.23.481644

**Authors:** Anna Z. Mykytyn, Melanie Rissmann, Adinda Kok, Miruna E. Rosu, Debby Schipper, Tim I. Breugem, Petra B. van den Doel, Felicity Chandler, Theo Bestebroer, Maurice de Wit, Martin E. van Royen, Richard Molenkamp, Bas B. Oude Munnink, Rory D. de Vries, Corine GeurtsvanKessel, Derek J. Smith, Marion P. G. Koopmans, Barry Rockx, Mart M. Lamers, Ron Fouchier, Bart L. Haagmans

**Author notes:** These authors contributed equally to this work (Melanie Rissmann, Adinda Kok, Miruna E. Rosu).

## Abstract

The emergence and rapid spread of SARS-CoV-2 variants may impact vaccine efficacy significantly^1^. The Omicron variant termed BA.2, which differs genetically substantially from BA.1, is currently replacing BA.1 in several countries, but its antigenic characteristics have not yet been assessed^2,3^. Here, we used antigenic cartography to quantify and visualize antigenic differences between SARS-CoV-2 variants using hamster sera obtained after primary infection. Whereas early variants are antigenically similar, clustering relatively close to each other in antigenic space, Omicron BA.1 and BA.2 have evolved as two distinct antigenic outliers. Our data show that BA.1 and BA.2 both escape (vaccine-induced) antibody responses as a result of different antigenic characteristics. Close monitoring of the antigenic changes of SARS-CoV-2 using antigenic cartography can be helpful in the selection of future vaccine strains.

---

Since its emergence in Wuhan, China, in 2019, SARS Coronavirus 2 (SARS-CoV-2) has caused over 300 million cases and 5.5 million deaths^4^. The initial virus that spread globally was characterized by a D614G change in the spike (S) protein^5^.Approximately one year into the pandemic other variants with 6-12 mutations in the S protein started to become dominant in various countries^6^. These variants included the Alpha variant in the United Kingdom, the Beta variant in South Africa, and the Gamma variant in Brazil, of which Alpha became the dominant variant globally by early 2021. In the summer of 2021, the Delta variant emerged first in India, and replaced Alpha globally within several months^7-10^. Emerging variants are termed Variants of Concern (VOC) by the World Health Organization (WHO) if they are associated with a major change in epidemiology and/or clinical presentation, increased virulence, increased transmissibility, and/or decreased effectiveness of public health and social measures or available diagnostics, vaccines or therapeutics ^11^. In addition, the WHO has designated other variants as Variants of Interest (VOI), which possess mutations predicted or known to affect antibody escape, virulence, or transmission. At the end of 2021 the VOC Omicron (sublineage BA.1) emerged in Botswana and South Africa, carrying 30 mutations in S, raising concerns for extensive immune evasion^1^. Whereas several earlier VOCs and VOIs exhibit some levels of antibody escape (e.g. Beta, Gamma, Delta, and Mu), they were still neutralized well by convalescent and post-vaccination sera^12-16^. In contrast, Omicron BA.1 almost completely escapes neutralizing antibodies, leading to low levels of remaining protective antibodies in most previously infected or vaccinated individuals, and a high frequency of breakthrough infections. This antibody escape at least partly explains why this variant has become the dominant variant globally over the span of a few weeks^17-20^. Fortunately, BA.1 appears to be less virulent compared with earlier variants^21,22^. A second Omicron variant (sublineage BA.2), emerged in South Africa around the same time as BA.1 and differs from BA.1 by 40 mutations (including 18 substitutions and 5 indels in S)^2^. Whereas BA.2 initially was only sporadically detected, it is currently replacing BA.1 in several countries^3^. It is unclear why BA.2 is spreading so fast, but it may be inherently more transmissible or considerably antigenically distinct from BA.1 and earlier variants.

As SARS-CoV-2 continues to evolve and escape neutralizing antibody responses, it is becoming increasingly important to understand the antigenic relationships among variants and the substitutions that underlie antigenic changes. Antigenic cartography is a tool to quantitatively analyze antigenic drift and visualize the emergence of new antigenic clusters, which is why it is used to biannually inform influenza virus vaccine strain selection^23,24^. Here, we investigated the neutralizing activity of human post-vaccination sera to both Omicron variants and applied antigenic cartography to 11 SARS-CoV-2 variants and hamster sera elicited against 8 SARS-CoV-2 variants by primary infection.

## Neutralizing activity of human sera against Omicron BA.1 and BA.2

Multiple studies have shown that Omicron BA.1 efficiently evades antibody responses post-infection and post-vaccination^12-16^. However, few studies have analyzed antibody responses to Omicron BA.2. Therefore, we investigated to what extent human post-BNT162b2 vaccination sera neutralized Omicron BA.2. After a single BNT162b2 vaccination, on average low neutralizing titers were detected against all variants with a 13 and 8-fold drop in neutralizing titers against Omicron BA.1 and BA.2, respectively, compared with 614G. In line with previous findings, Omicron BA.1 was 61-fold less efficiently neutralized after two BNT162b2 vaccinations^18-20^ (Fig. 1a-b). In comparison, Omicron BA.2 was neutralized somewhat more efficiently with a 13-fold drop in neutralizing titers compared with 614G. A third vaccination with BNT162b2 reduced the fold change to 614G to 11 and 7-fold for BA.1 and BA.2, respectively. Titers against all variants increased with each dose (Fig. 1a-c). Combined, these data show that Omicron BA.1 and BA.2 escape antibody responses, and suggest that the height and breadth of the antibody response against SARS-CoV-2 can be increased by repeated exposure to the original antigen. The differential effect of booster vaccination on BA.1 and BA.2 suggested that both variants are antigenically distinct, warranting further analysis. Sera from primary infections with SARS-CoV-2 variants can be used to assess the antigenicity of different variants, however human sera from primary variant infections are increasingly difficult to obtain. We did attempt to obtain human sera post-primary Omicron infection and included 12 individuals who had not been vaccinated, of which 4 reported a previous SARS-CoV-2 infection. 6 had high neutralizing titers against 614G and Delta, suggesting that an additional 2 individuals had been infected with another variant prior to their BA.1 infection (Extended Data Fig. 1a). The remaining 6 sera had low neutralizing activity against all variants, including Omicron (geometric mean titer of 14, 32, 63 and 41 against 614D, Delta, Omicron BA.1 and BA.2 respectively) (Extended Data Fig. 1b). As primary antisera against future variants will become even more difficult to obtain, we determined the antigenicity of SARS-CoV-2 variants using the hamster model. Hamsters were used as they are highly susceptible to SARS-CoV-2 and are therefore ideal for controlled infections and obtaining well defined sera^25-27^.

**Fig. 1.**
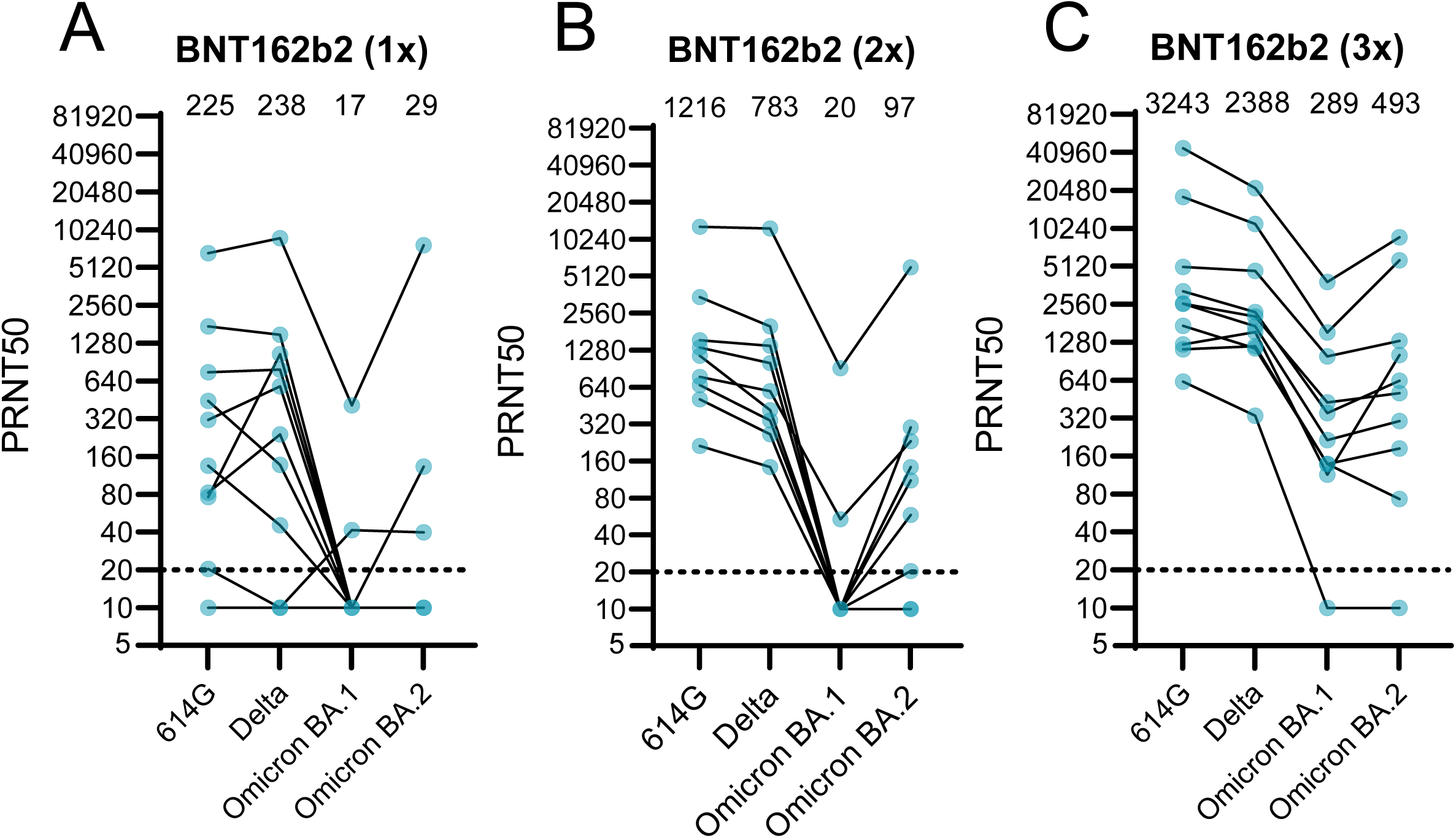
Neutralizing activity of human post-vaccination sera against Omicron BA.1 and BA.2. **a-c**, Neutralization titers against 614G, Delta, Omicron BA.1 and Omicron BA.2 of vaccinated individuals after vaccination with one (**a**), two (**b**) or three (**c**) doses of BNT162b2. Geometric mean is displayed above each graph. PRNT50: plaque reduction neutralization titers resulting in 50% plaque reduction. Dotted lines indicate limit of detection.

### Antigenic cartography of SARS-CoV-2

To investigate the antigenic relationships between BA.1, BA.2 and other SARS-CoV-2 variants, we used antigenic cartography^28^. We used the Syrian golden hamster model to generate antisera by inoculating hamsters with SARS-CoV-2 variants (614G, Alpha, Beta, Gamma, Zeta, Delta, Mu and Omicron (lineage BA.1)) (Fig. 2a, Extended Data Fig. 2). Virus stocks and the original respiratory specimens were sequenced to confirm the absence of culture-acquired mutations that may affect antigenic properties. In addition, at 7 days post-infection (dpi) swabs were collected and sequenced to confirm that the virus did not change during the experiment. Apart from Delta and Omicron BA.1-infected animals, whose swabs yielded too little virus to sequence, no mutations in Spike were detected in swab sequences. Plaque reduction neutralization titers resulting in a 50% reduction in infected cells (PRNT50) obtained with both authentic SARS-CoV-2 and pseudotyped viruses were used to generate antigenic maps (Fig. 2b). All animals were successfully infected, as shown by high viral RNA titers at 1 dpi in nasal washes and high homologous antibody titers at 26 dpi (Fig. 2c-d).

**Fig. 2.**
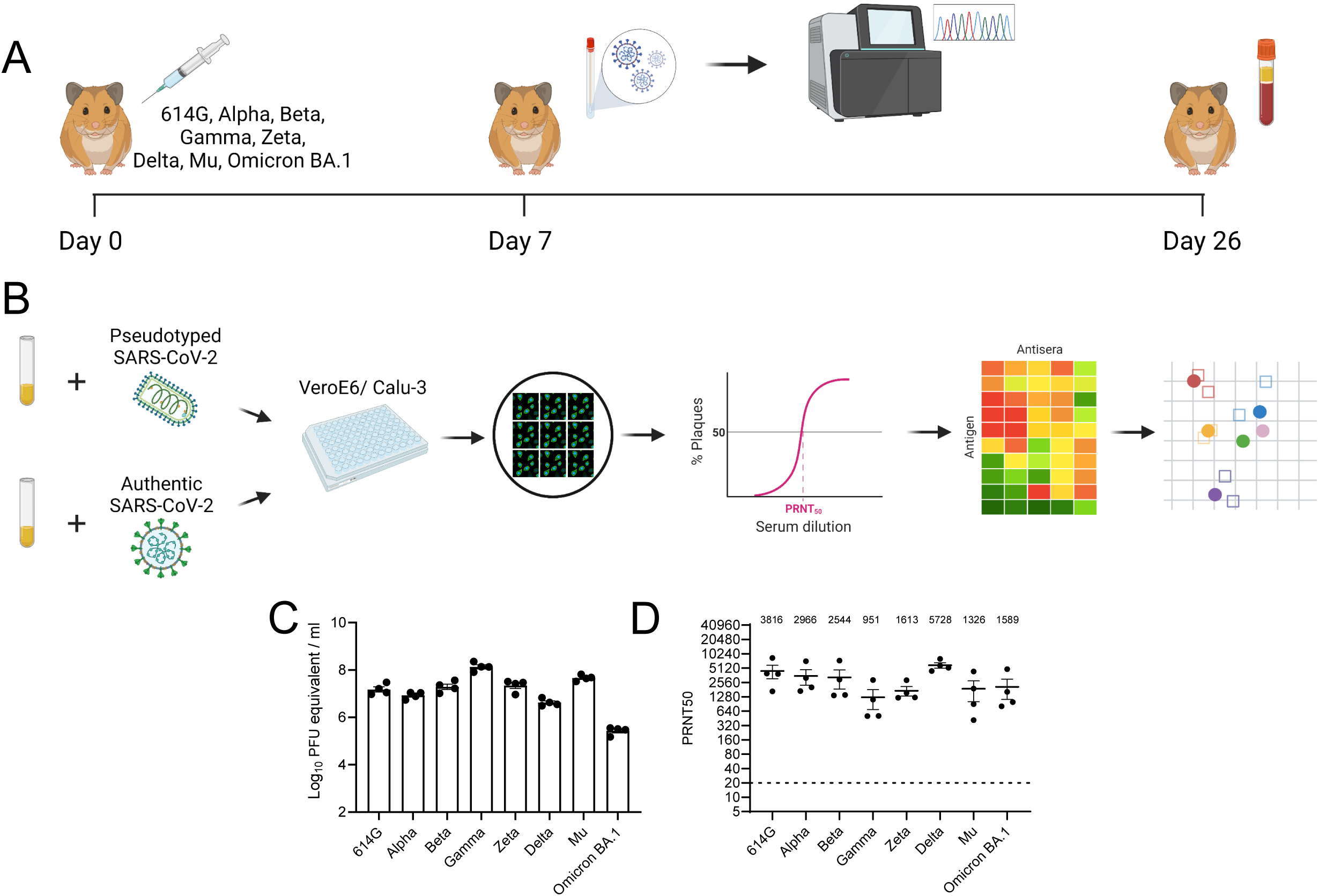
SARS-CoV-2 variants efficiently infect hamsters, inducing high neutralizing antibody titers. **a**, Hamsters were inoculated with the indicated SARS-CoV-2 variants. Nasal washes were collected 7 days post-infection and sequenced. At 26 days post-infection blood was collected for serological analysis. **b**, Hamster sera was assessed for neutralizing antibodies against pseudotyped and authentic SARS-CoV-2. PRNT50 values were used to generate antigenic maps using a multidimensional scaling algorithm. **c**, RNA titers of nasal washes collected one day post-infection. **d**, Homologous PRNT50 titers were determined using authentic SARS-CoV-2. Geometric mean is displayed above graph. PRNT50: plaque reduction neutralization titers resulting in 50% plaque reduction. Dotted line indicates limit of detection. Error bars indicate SEM. Panels **a** and **b** were created with BioRender.com.

Pseudovirus neutralization assays are a safe and widely used tool to assess antibody neutralization. We performed initial neutralization experiments on VeroE6 cells, which are the most commonly used cell line for neutralization assays. We used the spike variants 614D, 614G, Alpha, Beta, Delta, Kappa and Omicron BA.1. All homologous sera neutralized to high titers (Extended Data Fig. 3a-h). The lowest cross-neutralizing titers were obtained against Omicron for all sera with 9 to 43-fold reduction compared to homologous neutralization. Sera from Omicron BA.1-infected hamsters poorly neutralized all other variants (2 to 81 fold reduction compared to homologous neutralization). These data show that Omicron BA.1 induces different antibody responses compared with all other variants. Next, we performed pseudovirus neutralization assays on the human airway Calu-3 cell line. In Calu-3 cells SARS-CoV-2 enters using the serine protease-mediated entry pathway that is also used in primary cells, whereas in VeroE6 cells SARS-CoV-2 enters using the cathepsin-mediated endocytic entry pathway^29^. In addition, the variability in infectivity between variants is lower for Calu-3 cells compared with VeroE6, suggesting that Calu-3 cells may allow for more equal comparisons^30^. Neutralization titers on Calu-3 cells were similar to VeroE6 and correlated well (Extended Data Fig. 3i-p, Extended Data Fig. 4a-g).

Next, we constructed antigenic maps from the neutralization data using a multidimensional scaling (MDS) algorithm described previously^28^ (Fig. 3a-b). These were verified to ensure fitted distances correlated well with actual neutralizing titers and to confirm that the data was well represented in two dimensions (Extended Data Fig. 5a-d). We found that the map constructed by Calu-3 cells was very similar to the VeroE6 map, since the same antigens plotted within one 2-fold dilution from each other in the two maps (Extended Data Fig. 6). Therefore, the choice of the cell line for the neutralization assay did not affect the topology of the map substantially. In order to assess whether the map generated with pseudovirus neutralization data would accurately represent authentic SARS-CoV-2, we generated a map with the same antisera as in Fig 3a-b and 614G, Alpha, Beta, Delta and Omicron viruses on Calu-3 cells. (Fig. 3c). We confirmed that there was a good correlation between the raw neutralizing titers of the 5 variants on Calu-3 and VeroE6 cells (Extended Data Fig. 7-8). The map generated with authentic SARS-CoV-2 closely resembled the maps generated with pseudovirus, the positions of the antigens differed no more than one 2-fold dilution between maps (Fig. 3d). Antigenically, we found that all variants aside from Omicron BA.1 grouped closely together. Omicron BA.1 formed a distinct antigenic outlier in the map, 10 to 38-fold dilutions away from the nearest virus (Fig. 3a-c).

**Fig. 3.**
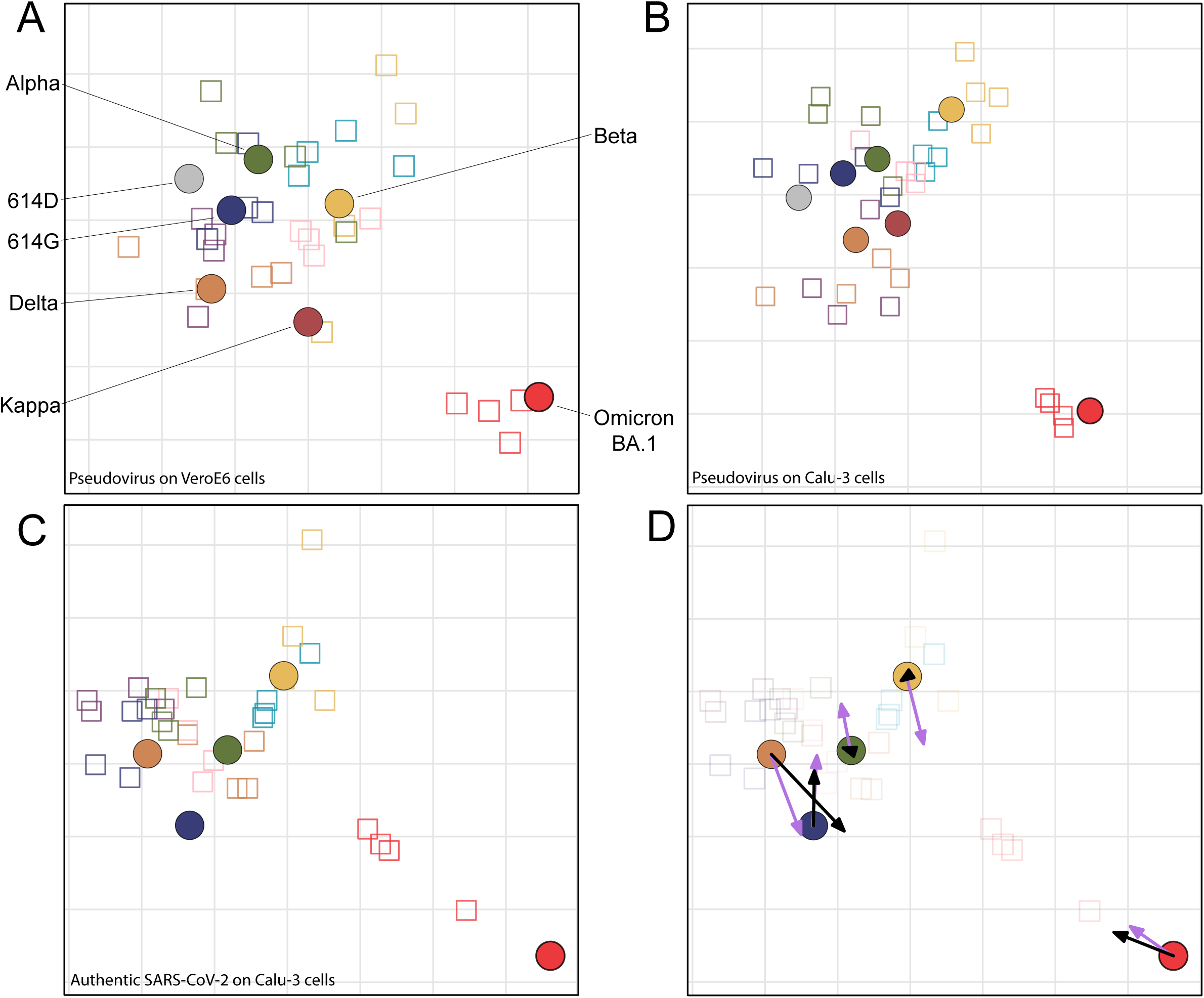
Antigenic maps comparing neutralizations with SARS-CoV-2 pseudoviruses and authentic SARS-CoV-2. **a-b**, MDS was used to create an antigenic map from the PRNT50 titers generated against 614D, 614G, Alpha, Beta, Delta, Kappa and Omicron pseudoviruses on either VeroE6 (**a**) or Calu-3 (**c**) cells. **c**, MDS was used to create an antigenic map from the PRNT50 titers generated against 614G, Alpha, Beta, Delta and Omicron authentic SARS-CoV-2. **d**, Re-display of antigenic map in **c** with lilac arrows indicating antigen positions in map **a** and black arrows indicating antigen positions in map **b**. Viruses are shown as coloured circles and antisera as squares with the same outline colour as the matching viruses. Viruses and antisera are positioned in the map so that the distances between them are inversely related to the antibody titers, with minimized error. The grid in the background scales to a 2-fold dilution of antisera in the titrations. MDS: multidimensional scaling. PRNT50: plaque reduction neutralization titers resulting in 50% plaque reduction.

Next, we extended the authentic virus dataset to contain a larger set of variants: 614G, Alpha, Beta, Gamma, Zeta, Delta, Delta AY.4.2, Lambda, Mu, Omicron BA.1 and Omicron BA.2. As for pseudotyped virus, homologous sera neutralized to a high titer across all variants (Fig. 4a-h, homologous titers per panel are a re-display from Fig. 1d). Similar to pseudovirus data, we observed a reduction in neutralization titers of Omicron BA.1 sera against all other variants (2.4 to 9 fold compared to homologous) and poor neutralization of Omicron BA.1 by all non-homologous sera (8 to 112 fold reduction). In addition, Omicron BA.2 was also poorly neutralized by all sera (7 to 114 fold reduction), including Omicron BA.1 (8 fold reduction). Although Omicron BA.1 and Omicron BA.2 possess many overlapping mutations in S, the differences between the variants were sufficient to prevent efficient cross-neutralization. In agreement, our study shows that antibodies elicited against the original SARS-CoV-2 cluster do not neutralize Omicron BA.1 well, and vice-versa.

**Fig. 4.**
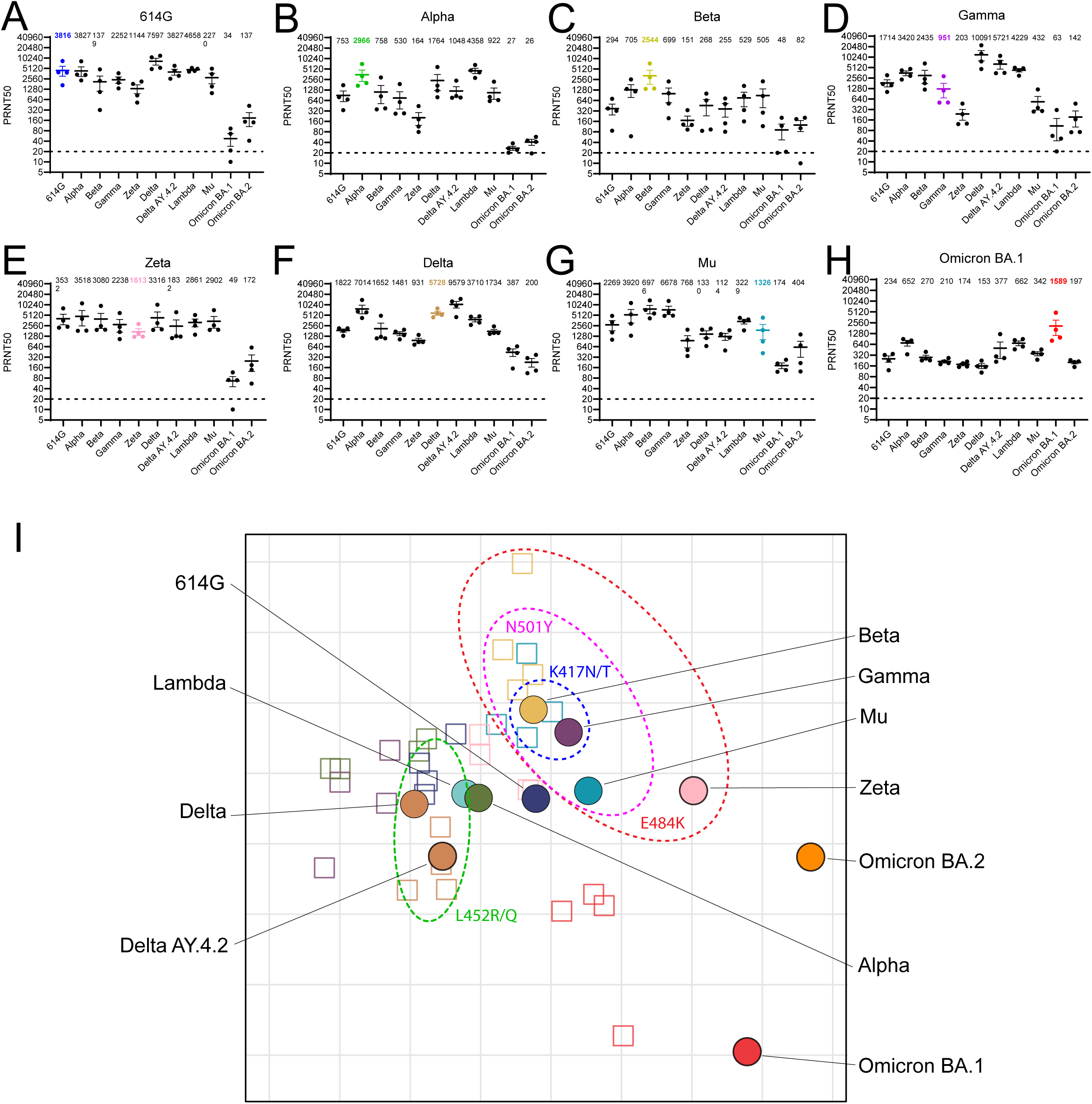
Antigenic cartography using authentic SARS-CoV-2. **a-h**, Neutralizing titers of hamsters infected with either (**a**) 614G, (**b**) Alpha, (**c**) Beta, (**d**) Gamma, (**e)** Zeta, (**f**) Delta, (**g**) Mu or (**h**) Omicron BA.1 viruses. **i**, Multidimensional scaling was used to create an antigenic map utilizing PRNT50 titers generated from authentic SARS-CoV-2 on Calu-3 cells. See legend to Fig. 3 for details. Subdivided by dotted ellipses are variants possessing overlapping substitutions as indicated. Geometric mean is displayed above each graph. PRNT50: plaque reduction neutralization titers resulting in 50% plaque reduction. Dotted lines indicate limits of detection. Error bars indicate SEM.

The generated antigenic map based on the extended neutralization dataset was verified by ensuring fitted distances correlated well with actual neutralizing titers, that the data was well represented in two dimensions and plotted the uncertainty of each antigen and antisera (Extended Data Fig. 9). The antigenic map shows that, similarly to the maps in Fig. 3, all variants except Omicron BA.1 and BA.2 group together in one antigenic cluster (Fig. 4i). In line with the limited cross-neutralization of Omicron BA.2 with BA.1 sera, both variants were positioned distantly from each other in the map, with BA.2 somewhat closer to the main cluster than BA.1, indicating that Omicron BA.1 induced qualitatively different antibody responses and BA.1 and BA.2 are antigenically distinct SARS-CoV-2 variants. The antisera corresponding to each virus grouped in the same region of the map, indicating efficient homologous neutralization. As expected, based on Fig. 3, 614G and Alpha are in the center of the cluster. Within this cluster, viruses grouped together based on specific substitutions. Viruses containing E484K (Beta, Gamma, Zeta and Mu) grouped to the top-right of the ancestral 614G virus, whereas viruses containing substitutions L452R/Q grouped to the left of 614G. The Beta and Gamma variants, which in addition to E484K both contain K417N/T also cluster together. The same clustering based on E484K and L452R/Q was observed in the pseudovirus maps. A recently study by Wilks and colleagues used human sera to generate an antigenic map^31^. Both the maps from our study and the map from Wilks and colleagues show the same clustering of variants containing mutations at position 484, 417 and 452, indicating that hamsters and humans generate similar antibody responses. However, differences were observed as well. In the map by Wilks and colleagues, there was approximately a ∼2 fold larger distance between 614G and variants Beta, Gamma and Mu, compared to our map containing all authentic viruses. This may be caused by specific antigenic relationships between the large set of variants, as including only 5 variants increased the distance from 614G to Beta by ∼2 fold (Fig. 3c). In addition, these differences may be caused by the lower titers observed in naturally-infected humans compared with experimentally-infected hamsters. Low titers against immunizing viruses may drop off more for viruses with only a few immune-evasive substitutions (e.g. Beta, Gamma, and Delta), leading to relatively large antigenic distances. On the other hand, low titers may underestimate the antigenic distance for highly evasive viruses due to reaching the assay’s lower limit of detection. In agreement, the distance of the main cluster to Omicron BA.1 was larger in our map compared with the map by Wilks and colleagues. Nevertheless, these data indicate that human and hamster serum responses to SARS-CoV-2 infection are similar as they lead to topologically similar maps. However, the specific set of viruses and sera used to construct an antigenic map may influence antigenic distances, e.g. due to omission of sera against Omicron. As human sera post-primary infection are increasingly difficult to obtain, our data suggest that antigenic cartography using hamster sera is a useful surrogate to assess antigenic relationships between SARS-CoV-2 variants.

The emergence and rapid spread of the heavily mutated Omicron BA.1 and BA.2 variants suggests that population immunity is exerting strong selective pressure on SARS-CoV-2, favoring the emergence of new antigenic variants. As the number of SARS-CoV-2 variants increases it will become increasingly important to visualize and understand the antigenic relationships between variants. Here, we used antigenic cartography to quantify and visualize SARS-CoV-2 antigenic evolution and demonstrate that Omicron BA.1 and BA.2 have evolved as two antigenically distinct variants, separate from an ancestral cluster with all earlier SARS-CoV-2 variants.

The evolutionary history of SARS-CoV-2 in humans is relatively short compared with viruses that have circulated in humans for decades, such as influenza viruses^32^. Before the emergence of Omicron, most SARS-CoV-2 variants contained only few substitutions in the S protein and were still recognized by convalescent and post-vaccination sera. These variants all grouped into the same antigenic cluster in our antigenic maps, but within that cluster a grouping could be observed based on S substitutions at position 417, 484 and 452, in line with previous data on human sera^31^. The Omicron variants, however, positioned as distinct antigenic variants with limited cross-neutralization to each other and the original cluster. These data were corroborated by human post-vaccination antibody responses (Fig. 1). The ameliorating effect of the booster vaccine on Omicron BA.2 responses suggests that population immunity against BA.2 may be sufficient to prevent flooding of healthcare systems and high levels of mortality, seen before for previous variants^33-35^. In addition, the widespread circulation of BA.1 may lead to the broadening of the antibody response in previously infected or vaccinated individuals and subsequent dampening of the intensity of spread. However, in regions with low access to vaccines, a wave of primary infections with BA1 would potentially lead to low level cross-protection and continued opportunity for further widespread circulation of BA2 or other variants.

The antigenic cartography of SARS-CoV-2 visualizes clearly how BA.1 and BA.2 can both escape antibody responses without being antigenically similar. The emergence of both Omicron variants indicates that population immunity is selecting for SARS-CoV-2 variants that efficiently escape from neutralizing antibody responses, leading to the first signs of antigenic drift. SARS-CoV-2 will eventually reach endemicity and likely cause annual or biannual infection waves as seen for influenza and seasonal coronaviruses. Our study lays the foundation for the continuous monitoring of the antigenic evolution of SARS-CoV-2, which may inform the selection of vaccine strains in the future.

## Supplementary methods

### Viruses and cell lines

HEK-293T cells were cultured in Dulbecco’s modified Eagle’s medium (DMEM) supplemented with 10% FCS, sodium pyruvate (1 mM, Gibco), non-essential amino acids (1x, Lonza), penicillin (100 IU/ mL), and streptomycin (100 IU/mL). VeroE6 cells were maintained in DMEM supplemented with 10% FCS, HEPES (20 mM, Lonza) and sodium pyruvate (1 mM), penicillin (100 IU/mL), and streptomycin (100 IU/mL). Calu-3 cells were maintained in Eagle’s minimal essential medium (ATCC) supplemented with 10% FCS, penicillin (100 IU/mL), and streptomycin (100 IU/mL). All cell lines were kept at 37°C in a humidified CO_2_ incubator. Viruses propagated for infection in hamsters and neutralization assays were grown to passage 3 on Calu-3 cells (aside from 614G used in hamster inoculations, grown to passage 3 on Vero cells), harvested 48-72 hours post-infection, cleared for 5 minutes at 1000 x g, aliquoted and stored at -80°C until use. All work with infectious SARS-CoV-2 was performed in a Class II Biosafety Cabinet under BSL-3 conditions at Erasmus Medical Center.

### Pseudovirus production

Pseudoviruses were produced as described previously^29^. Briefly, HEK-293T cells were transfected with pseudotyping vectors from InvivoGen (Original D614, D614G, Alpha, Beta, Kappa and Delta Spike) or kindly provided by Dr. Berend Jan Bosch (Omicron Spike). All Spike expressing plasmids contained a deletion of the last 19 amino acids containing the Golgi retention signal of the SARS-CoV-2 Spike protein. Twenty four hours post-infection cells were infected with pseudoviruses expressing VSV-G. Two hours posti-infection, cells were washed three times with Opti-MEM I (1×) + GlutaMAX and replaced with medium containing anti-VSV-G neutralizing antibody (Absolute Antibody). The supernatant was collected after 24 and 48 hr, cleared by centrifugation at 2000 x g for 5 min, and stored at 4°C.

### Virus titrations

Stock titers were determined by preparing 10-fold serial dilutions in Opti-MEM I (1X) + GlutaMAX. One hundred ml of each dilution was added to monolayers of Calu-3 cells in the same medium in a 12 well plate. After 4 hours at 37°C, cells were overlaid with 1.2% Avicel (FMC biopolymers) in Opti-MEM I (1X) + GlutaMAX (Gibco) for 72 hr. Avicell was then removed and plates were fixed in formalin, permeabilized in 70% ethanol and washed in PBS. Cells were blocked in 3% BSA (bovine serum albumin; Sigma) in PBS, followed by rabbit anti-nucleocapsid (Sino biological; 1:2000) in PBS containing 0.1% BSA. Plates were washed in PBS then stained with goat anti-mouse Alexa Fluor 488 (Invitrogen; 1:4000) in PBS containing 0.1% BSA. Plates were then washed again in PBS and scanned on the Amersham Typhoon Biomolecular Imager (channel Cy2; resolution 10 mm; GE Healthcare). Eight hour titers of SARS-CoV-2 and pseudoviruses were determined by preparing 10-fold serial dilutions in Opti-MEM I (1X) + GlutaMAX. Thirty ml of each dilution was added to monolayers of Calu-3 or VeroE6 cells in the same medium in a 96 well plate. After 16 hours at 37°C, Pseudovirus infected plates were fixed in paraformaldehyde and washed in PBS. After 8 hours at 37°C SARS-CoV-2 infected cells were fixed in formalin, permeabilized in 70% ethanol and washed in PBS. Cells were blocked in 3% BSA (bovine serum albumin; Sigma) in PBS, followed by rabbit anti-nucleocapsid (Sino biological; 1:2000) in PBS containing 0.1% BSA. Plates were washed in PBS then stained with goat anti-mouse Alexa Fluor 488 (Invitrogen; 1:4000) in PBS containing 0.1% BSA. SARS-CoV-2 and Pseudovirus infected cells were next stained with Hoechst (ThermoFisher) and washed with PBS. Cells were imaged using the Opera phenix spinning disk confocal HCS system (Perkin Elmer) equipped with a 10x air objective (NA 0.3) and 405 nm and 488 nm solid state lasers. Hoechst and GFP were detected using 435-480 nm and 500-550 nm emission filters, respectively. Nine fields per well were imaged covering approximately 50% of the individual wells. The number of green fluorescent protein (GFP) positive/ Alexa Fluor 488 positive infected cells were quantified using the Harmony software (version 4.9, Perkin Elmer).

### Hamster infections

Female Syrian golden hamsters (*Mesocricetus auratus*; 6 weeks old; Janvier, France) were handled in an ABSL-3 biocontainment laboratory. Groups of animals (n=4) were inoculated intranasally with 1.0×10^5 PFU (614G, Alpha, Beta, Gamma, Zeta, Mu) or 5.0×10^4 PFU (Delta, Omicron) in a total volume of 100μl per animal. Nasal washes (250μL) were taken at 7 dpi. All animals were sacrificed 26 dpi. Research involving animals was conducted in compliance with the Dutch legislation for the protection of animals used for scientific purposes (2014, implementing EU Directive 2010/63) and other relevant regulations. The licensed establishment where this research was conducted (Erasmus MC) has an approved OLAW Assurance # A5051-01. Research was conducted under a project license from the Dutch competent authority and the study protocol (#17-4312) was approved by the institutional Animal Welfare Body. Animals were housed in groups of 2 animals in filter top cages (T3, Techniplast), in Class III isolators allowing social interactions, under controlled conditions of humidity, temperature and light (12-hour light/12-hour dark cycles). Food and water were available ad libitum. Animals were cared for and monitored (pre-and post-infection) daily by qualified personnel. All animals were allowed to acclimatize to husbandry for at least 7 days. For unbiased experiments, all animals were randomly assigned to experimental groups. The animals were anesthetized (3-5% isoflurane) for all invasive procedures. Hamsters were euthanized by cardiac puncture under isoflurane anesthesia and cervical dislocation.

### Viral RNA quantification using qRT-PCR

RNA extraction was performed as described previously^36^. Briefly, 60 μL of sample was lysed in 90 μL of MagnaPure LC Lysis buffer (Roche) followed by a 15 minute incubation with 50 μL Agencourt AMPure XP beads (Beckman Coulter). Beads were washed twice with 70% ethanol on a DynaMag-96 magnet (Invitrogen) and eluted in 50 μL diethylpyrocarbonate treated water. qRT-PCR was performed using primers targeting the E gene^37^ and comparing the Ct values to a standard curve derived from a virus stock titrated on Calu-3 cells.

### Plaque reduction neutralization assay

Post-vaccination and post-Omicron BA.1 infection sera was kindly obtained in the scope of the healthcare worker study performed at the Erasmus MC. This study was approved by the institutional review board of the Erasmus MC (medical ethical committee, MEC-2020-0264). PRNT50 assays were performed as described previously. Briefly, sera was heat inactivated for 30 min at 56°C. Sera were 3-fold serially diluted in 60 μL Opti-MEM I (1X) + GlutaMAX (Gibco). Four hundred PFU (based on 8 hr titrations) in 60 μL were added per well to a final volume of 120 μL and a serum dilution of 1:20 in the first well. Plates were incubated for 1 hr at 37°C. Next 100 μL of virus and serum mix was added to confluent monolayers of Calu-3 or VeroE6 cells. SARS-CoV-2 infected plates were incubated for 8 hr at 37°C before fixing in formalin and permeabilizing in ethanol. Plates were then washed in PBS and stained as described for virus titrations. Pseudovirus infected plates were incubated for 16 hr at 37°C before fixing in paraformaldehyde and washing in PBS. Nuclei were stained with Hoechst for 30 min. Cells were imaged using the Opera phenix spinning disk confocal HCS system and the number of GFP-positive/ Alexa Fluor 488 positive infected cells were quantified using the Harmony software as described above. The PRNT50 was calculated using Graphpad Prism 9 based on non-linear regression.

### Illumina sequencing

Deep sequencing was performed as described previously^16^. Briefly, RNA was extracted as described above followed by cDNA synthesis and PCR using the QIAseq® SARS-CoV-2 Primer Panel kit (Qiagen) according to the manufacturer’s protocol. Omicron samples were amplified with an additional 11 primers (as described by ARTIC V4.1 primer set). Amplicons were purified using 0.8x AMPure XP beads followed by converting 100ng of DNA to paired-end Illumina sequencing libraries using the KAPA HyperPlus library preparation kit (Roche). Samples were pooled and analyzed on an Illumina sequencer V3 MiSeq flowcell (2×300 cycles). The 614G virus used for hamster infections was cultured to passage 3 on Vero cells and its S protein did not contain additional mutations. All other variant were propagated to passage 3 on Calu-3 cells. For neutralization assays another passage 3 614G isolate (Bavpat-1) was used with an identical S amino acid sequence (European Virus Archive Global #026 V-03883) and another passage 3 Beta isolate with an identical S amino acid sequence. The 614G Bavpat-1 passage 3 sequence was identical to the passage 1 (kindly provided by Dr. Christian Drosten). The Alpha (B.1.1.7), Gamma (P.1), Delta (B.1.617.2), Delta AY.4.2 (B.1.617.2 AY.4.2), Lambda (C.37), Mu (B.1.621), Omicron BA.1 (B.1.1.529 BA.1) and Omicron BA.2 (B.1.1.529 BA.2) variant passage 3 sequences were identical to the original respiratory specimens. Low coverage regions in Spike were confirmed by Sanger sequencing. The Beta variant (B.1.351, used for hamster inoculations) passage 3 sequence contained one synonymous mutation compared to the original specimen: A26449C (Wuhan-Hu-1 position) in E. The Beta variant (B.1.351, used in neutralization assays) passage 3 sequence contained two mutations compared the original respiratory specimen: one synonymous mutation C13860T (Wuhan-Hu-1 position) in ORF1ab and a L71P change in the E gene (T26456C, Wuhan-Hu-1 position). The Spike changes of all variants compared to Wuhan-Hu-1 are indicated in Extended data Fig. 2. All isolate sequences were submitted to Genbank. Hamster nasal wash sample sequences were identical to the input viruses.

### Antigenic cartography

Antigenic map construction was performed as described previously^28^. Briefly, antigenic cartography is a method to quantify and visualize neutralization data. In an antigenic map, the distance between antiserum point S and antigen point A corresponds to the difference between the log2 of the maximum titer observed for antiserum S against any antigen and the log2 of the titer for antiserum S against antigen A. Thus, each titer in a cross-titration can be thought of as specifying a target distance for the points in an antigenic map. Modified multidimensional scaling methods are then used to arrange the antigen and antiserum points in an antigenic map to best satisfy the target distances specified by the neutralization data. The result is a map in which the distance between the points represents antigenic distance as measured by the neutralization assay in which the distances between antigens and antisera are inversely related to the log2 titer. Because antisera are tested against multiple antigens, and antigens tested against multiple antisera, many measurements can be used to determine the position of the antigen and antiserum in an antigenic map, thus improving the resolution of the data. The antigenic maps were computed with the Racmacs package (https://acorg.github.io/Racmacs/, version 1.1.18.) in R. A web-based version of the software is available from https://www.antigenic-cartography.org/. The maps were constructed using 1000 optimizations, with the minimum column basis parameter set to “none”.

## Acknowledgments

The present manuscript was part of the research program of the Netherlands Centre for One Health

## Author contributions

Conceptualization: A.Z.M, M.M.L, R.F, B.R, B.L.H

Formal analysis: A.Z.M, M.R, A.K, M.E.R, M.M.L

Funding acquisition: B.H, M.P.K, R.R.d.V

Investigation: A.Z.M, M.R, D.S, T.B, PvD, F.C, T.B, M.d.W, M.E.R

Resources: B.B.O.M, C.G, R.R.d.V

Supervision: M.M.L, R.F, B.R, B.L.H

Writing—original draft: A.Z.M, M.M.L, B.L.H

Writing—review and editing: all authors reviewed and edited the final version.

## Data and materials availability

The sequences generated in this study have been deposited in GenBank. All data needed to evaluate the conclusions in the paper are present in the paper or the Supplementary Materials.

## Figure Legends

**Extended Data Fig.1.**
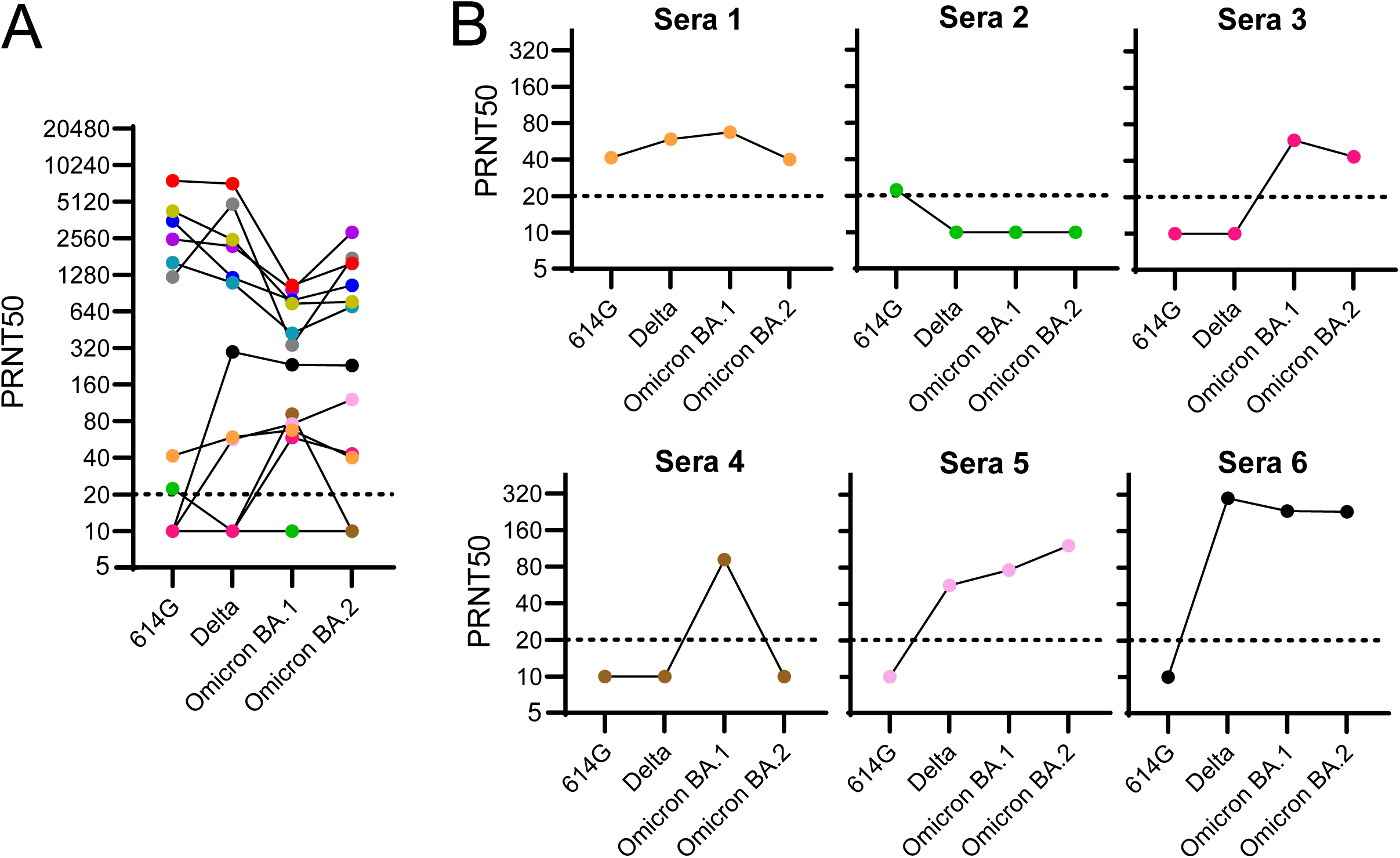
Neutralizing activity of human sera post Omicron BA.1 infection. **a**, Neutralization titers against 614G, Delta, Omicron BA.1 and Omicron BA.2 after primary Omicron BA.1 infection. **b**, Individual sera neutralization titers against 614G, Delta, Omicron BA.1 and Omicron BA.2 from 6 lowest responders in **a**. PRNT50: plaque reduction neutralization titers resulting in 50% plaque reduction. Dotted lines indicate limits of detection.

**Extended Data Fig.2.**
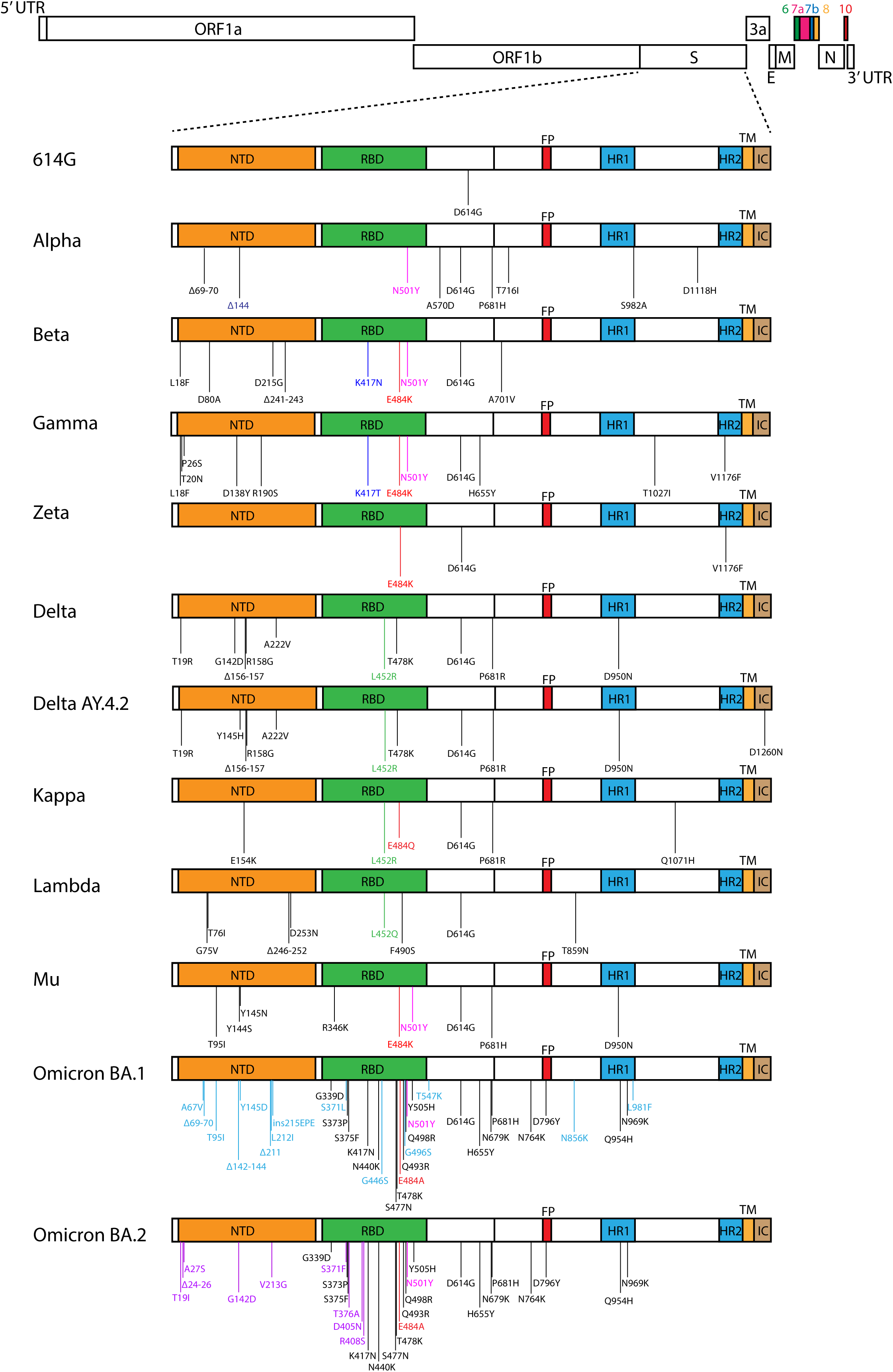
Spike substitutions in SARS-CoV-2 variants. Indicated are Spike substitutions present in all variants assessed. Indicated in red are substitutions at the 484 position, in pink substitutions at the 501 position and in dark blue substitutions at the 417 position. Differential substitutions between Omicron BA.1 and Omicron BA.2 are indicated in light blue and purple respectively.

**Extended Data Fig.3.**
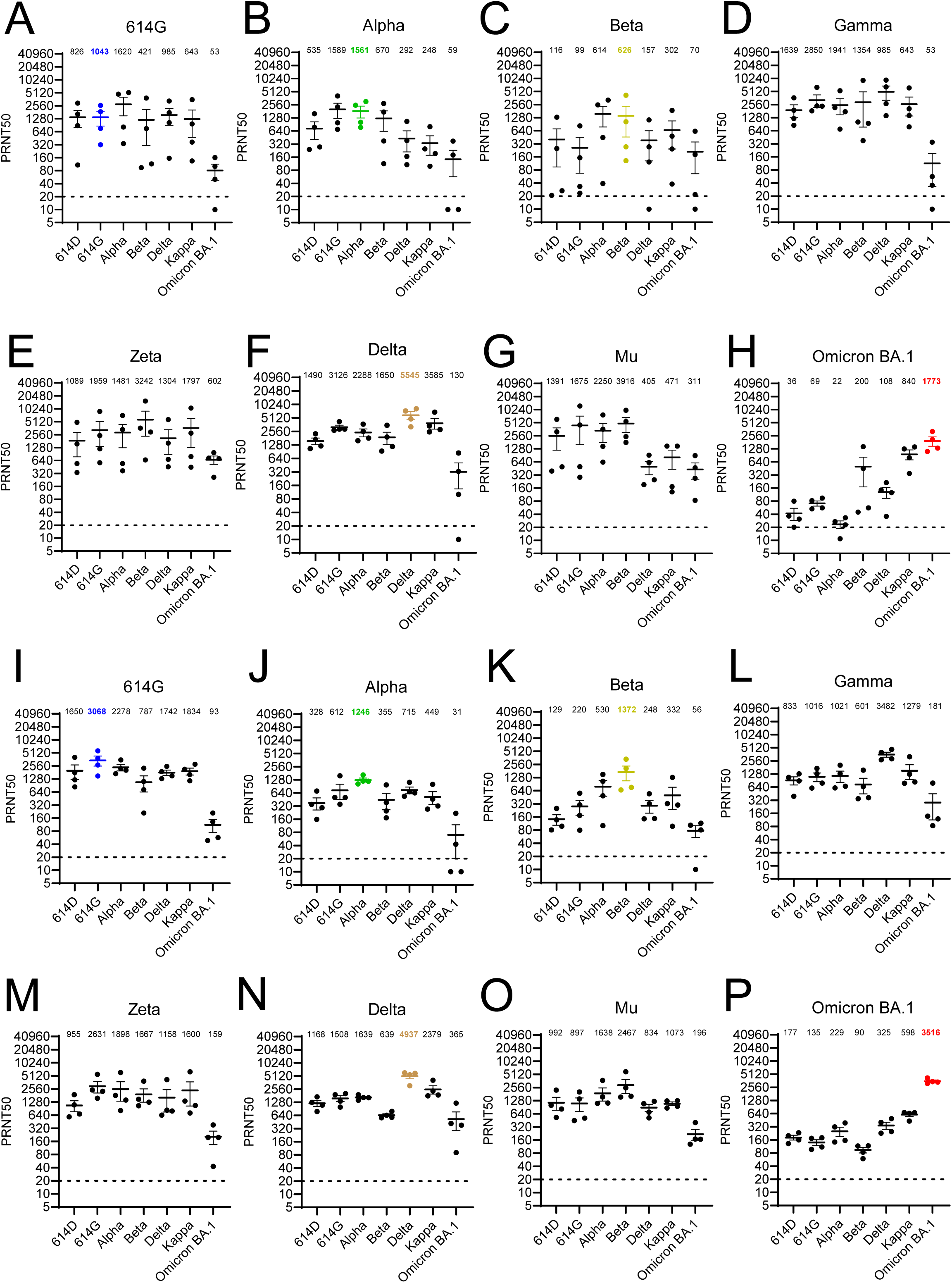
Pseudotyped SARS-CoV-2 neutralization titers. **a-h**, Neutralizing titers against pseudotyped SARS-CoV-2 on VeroE6 cells of hamsters infected with either (**a**) 614G, (**b**) Alpha, (**c**) Beta, (**d**)Gamma, (**e**) Zeta, (**f**) Delta, (**g**) Mu or (**h**) Omicron BA.1. i-p, Neutralizing titers against pseudotyped SARS-CoV-2 on Calu-3 cells of hamsters infected with either (**i**) 614G, (**j**) Alpha, (**k**) Beta, (**l**) Gamma, (**m**) Zeta, (**n**) Delta, (**o**)Mu or (**p**) Omicron BA.1. Geometric mean is displayed above each graph. PRNT50: plaque reductionneutralization titers resulting in 50% plaque reduction. Dotted lines indicate limits of detection. Error bars indicate SEM.

**Extended Data Fig.4.**
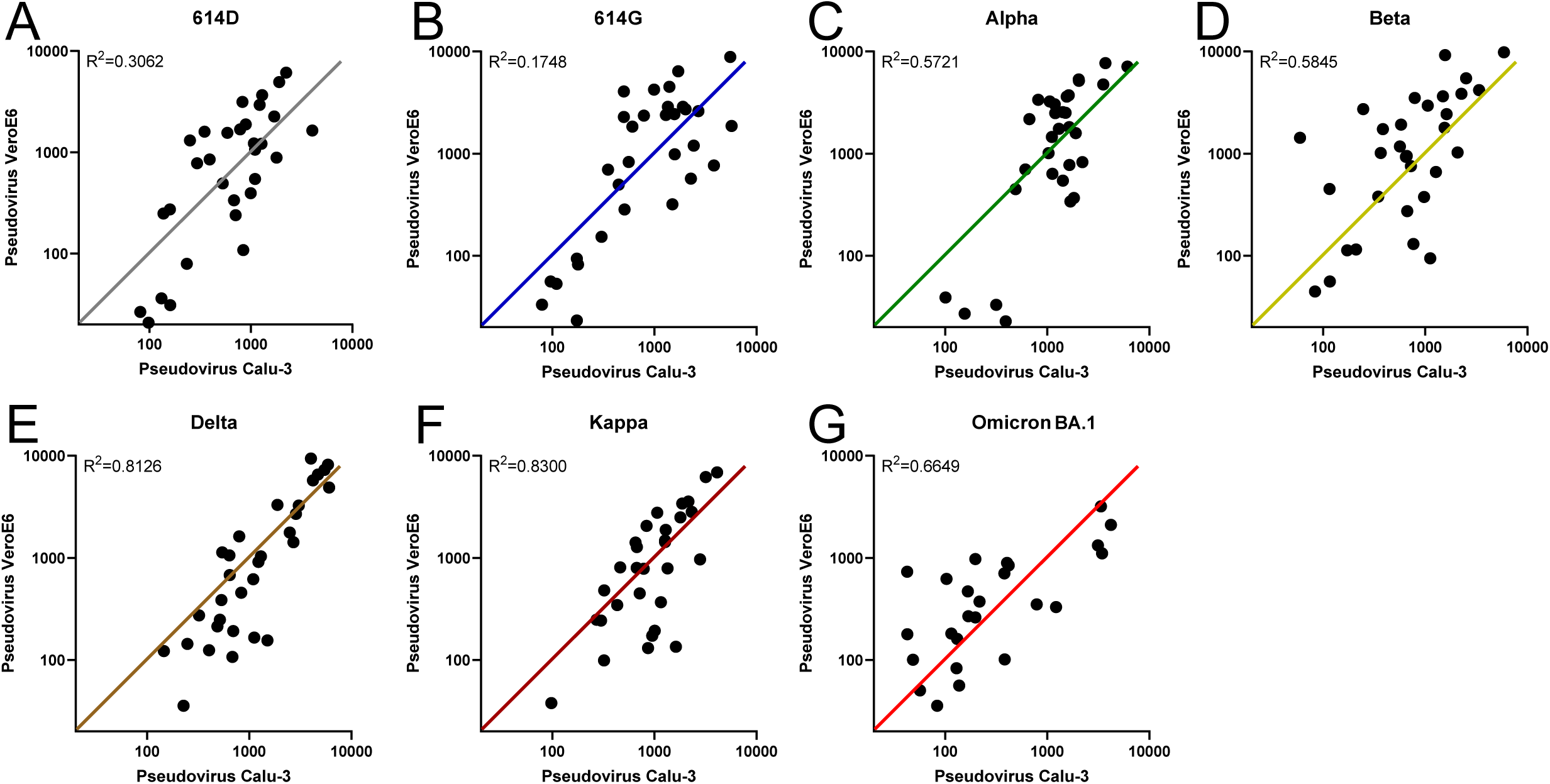
SARS-CoV-2 pseudovirus neutralization titers generated on VeroE6 cells correlate with titers generated on Calu-3 cells. **a-e**, Correlation of PRNT50 titers obtained with pseudotyped (**a**) 614D, (**b**) 614G, (**c**) Alpha, (**d**) Beta, (**e)** Delta, (**f**) Kappa and (**g**) Omicron BA.1 variants on VeroE6 cells compared to Calu-3 cells. PRNT50: plaque reduction neutralization titers resulting in 50% plaque reduction.

**Extended Data Fig.5.**
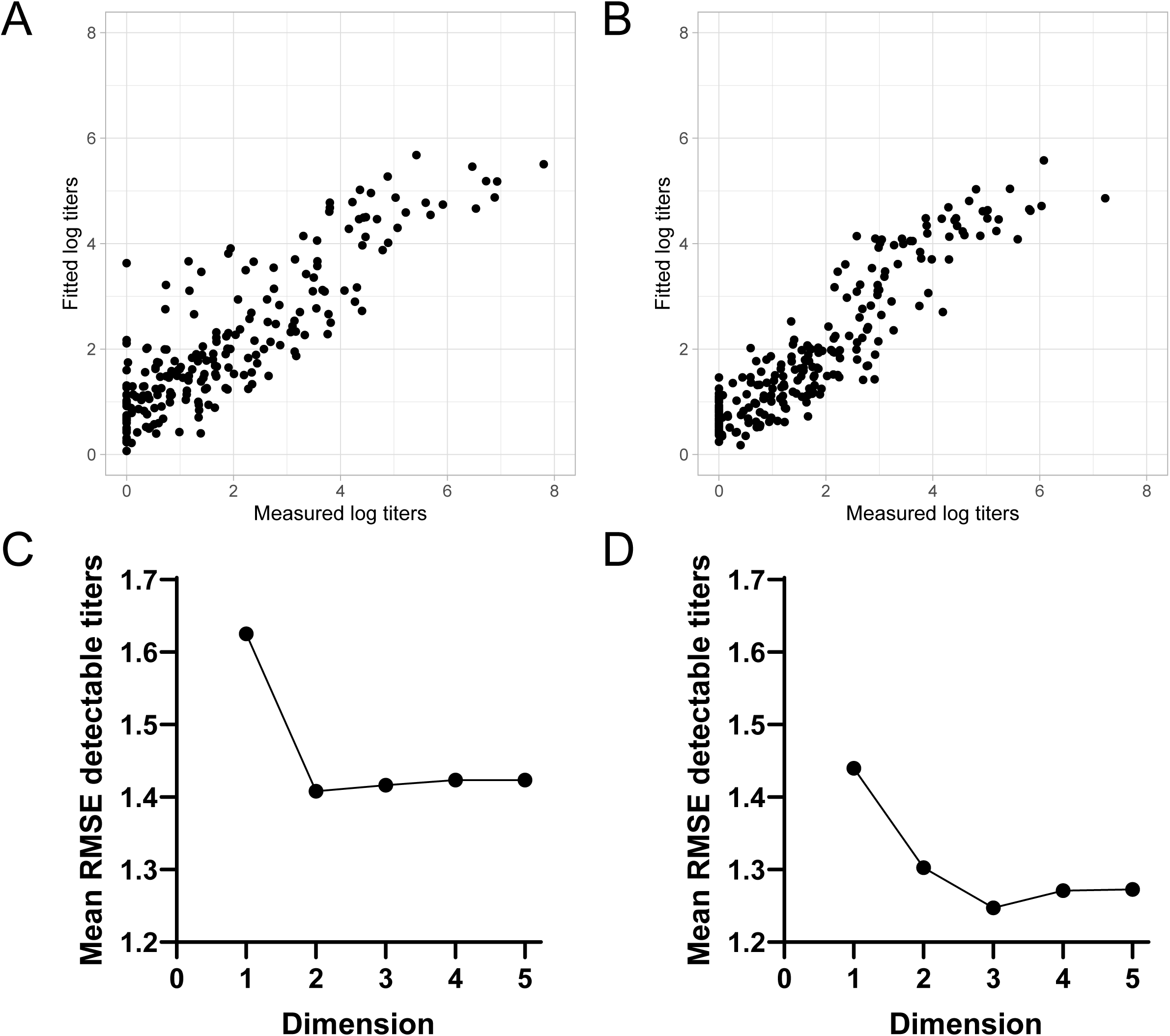
Validation of representing neutralization data in two dimensional antigenic maps. **a-b**, Scatter plot of detectable neutralizing titers on the x-axis and fitted titers from the antigenic maps on the y-axis of maps generated in Fig. 3 and 4. Scatter plots generated on data obtained with pseudovirus neutralizations on (**a**) VeroE6 cells (Fig. 3a) and (**b**) Calu-3 cells. **c-d**, Dimensionality tests indicating the mean RMSE of detectable neutralization titers in 1 to 5 dimensions for maps generated with data from neutralization on (**c**) VeroE6 cells or (**d**) Calu-3 cells. For each dimension 100 antigenic maps with 1000 optimizations each are generated while randomly excluding 10% of the titers. The mean RMSE is calculated by comparing the predicted titers in each run to the actual measured titers on a log2 scale. RMSE: root mean square prediction error.

**Extended Data Fig.6.**
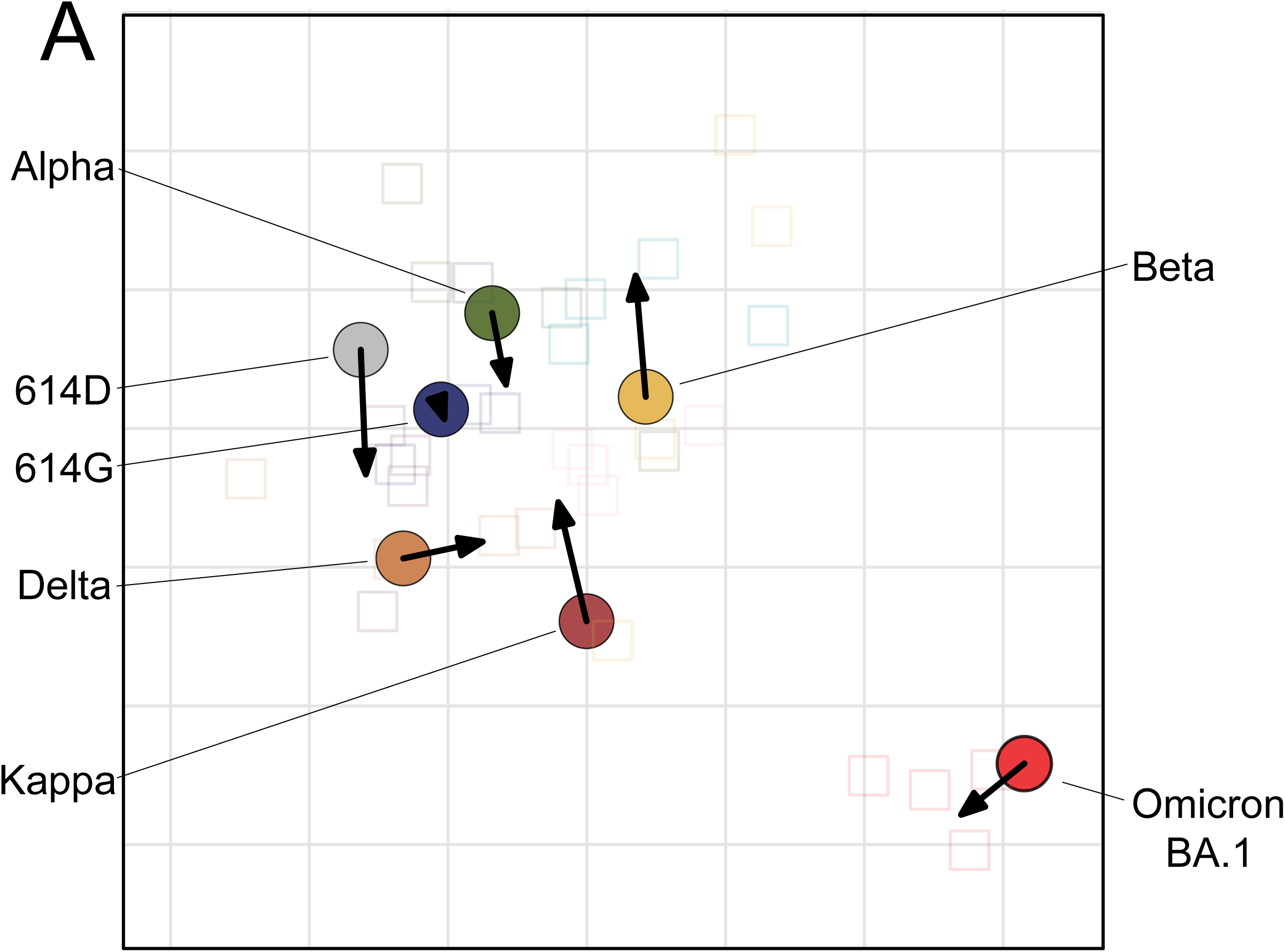
Comparison of the antigenic map generated with pseudovirus on VeroE6 cells to Calu-3 cells. Antigenic map from Fig. 3a with arrows indicating position of antigens in antigenic map in Fig. 3b. See legend to Fig. 3 for details.

**Extended Data Fig.7.**
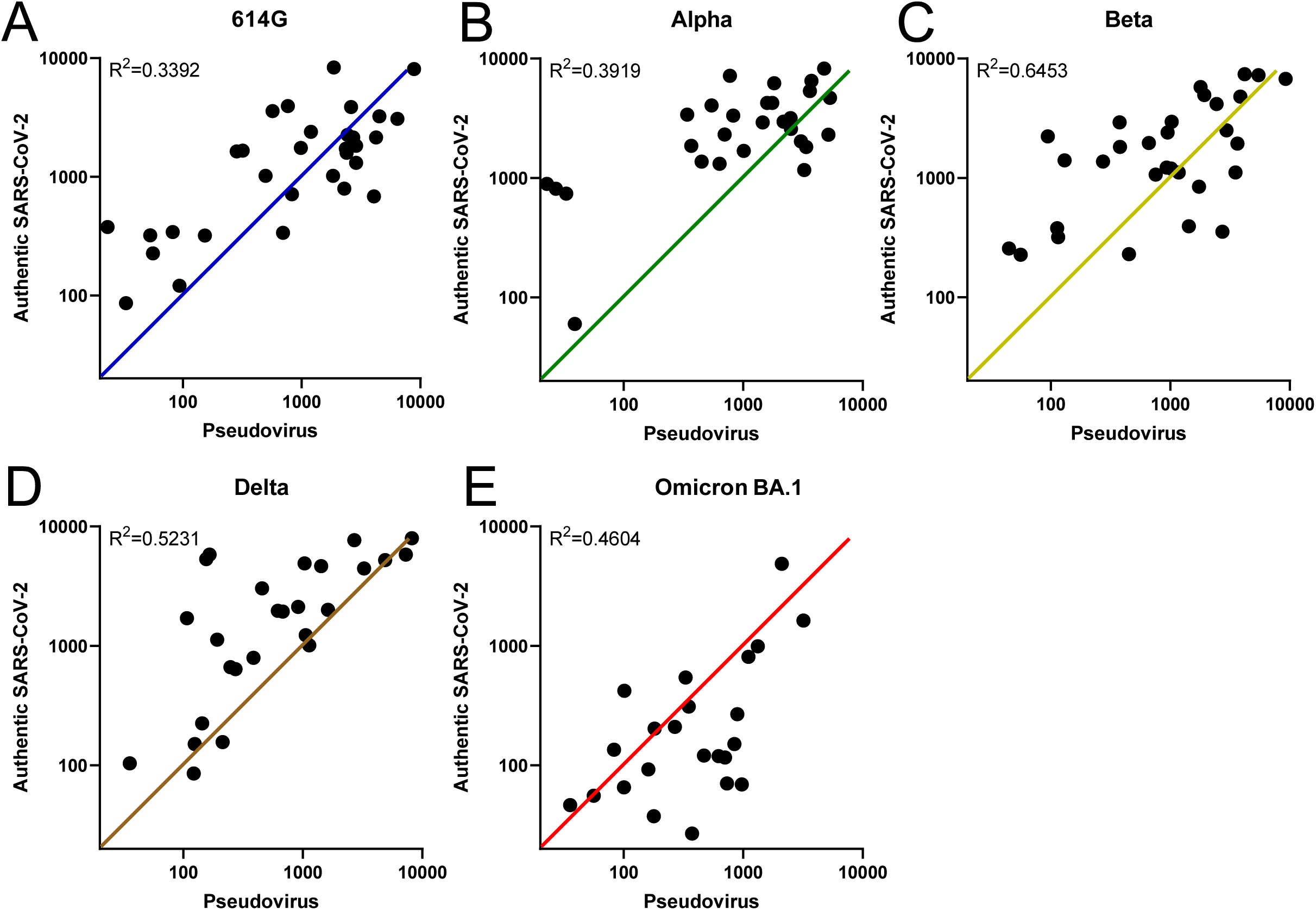
SARS-CoV-2 pseudovirus neutralization titers on VeroE6 cells correlate with authentic SARS-CoV-2 neutralization titers. **a-e**, Correlation of PRNT50 titers obtained with pseudotyped and authentic SARS-CoV-2 (**a**) 614G, (**b**) Alpha, (**c**) Beta, (**d**) Delta and (**e)** Omicron variants on VeroE6 cells. PRNT50: plaque reduction neutralization titers resulting in 50% plaque reduction.

**Extended Data Fig.8.**
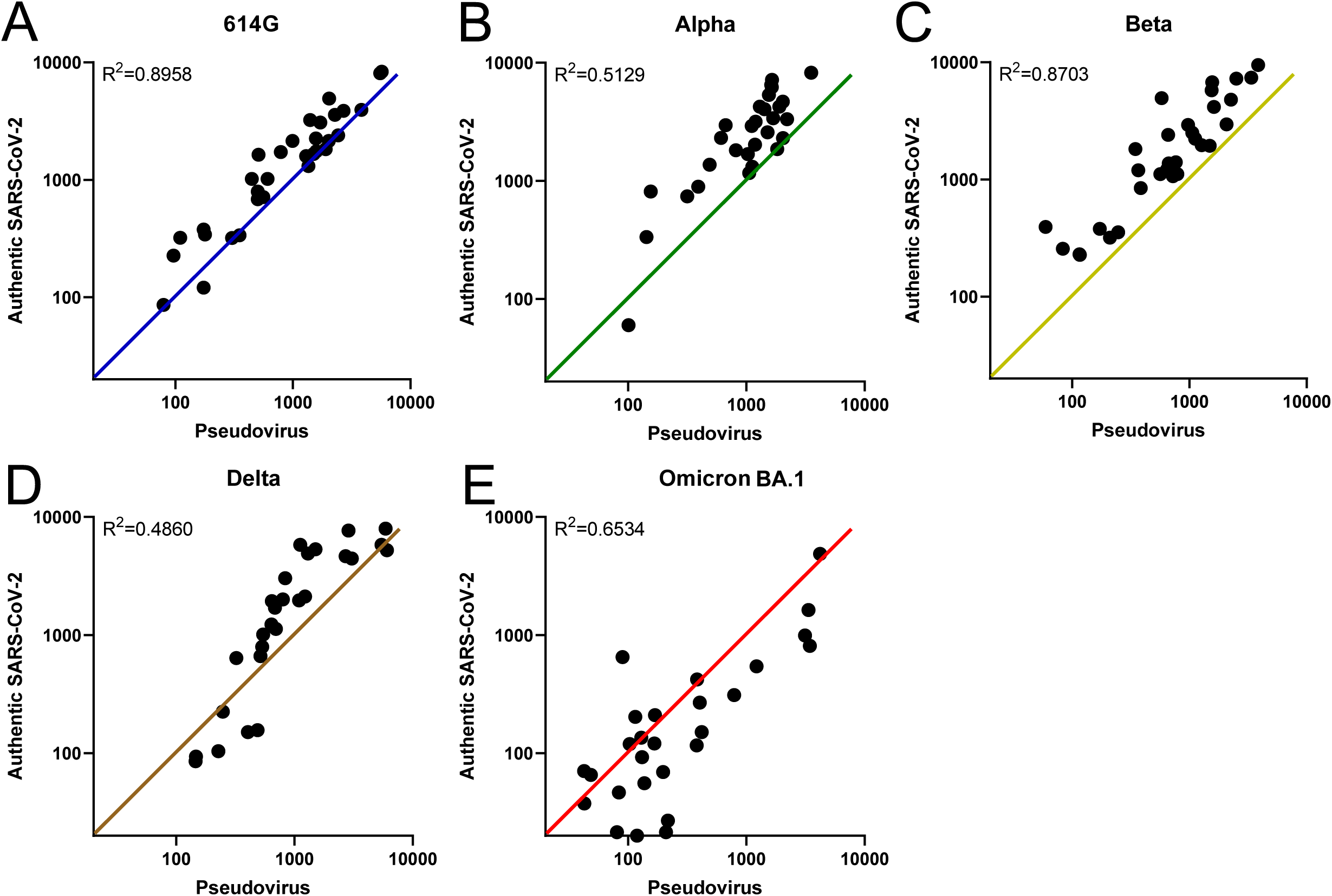
SARS-CoV-2 pseudovirus neutralization titers on Calu-3 cells correlate with authentic SARS-CoV-2 neutralization titers. **a-e**, Correlation of PRNT50 titers obtained with pseudotyped and authentic SARS-CoV-2 (**a**) 614G, (**b**) Alpha, (**c**) Beta, (**d**) Delta and (**e)** Omicron variants on Calu-3 cells. PRNT50: plaque reduction neutralization titers resulting in 50% plaque reduction.

**Extended Data Fig.9.**
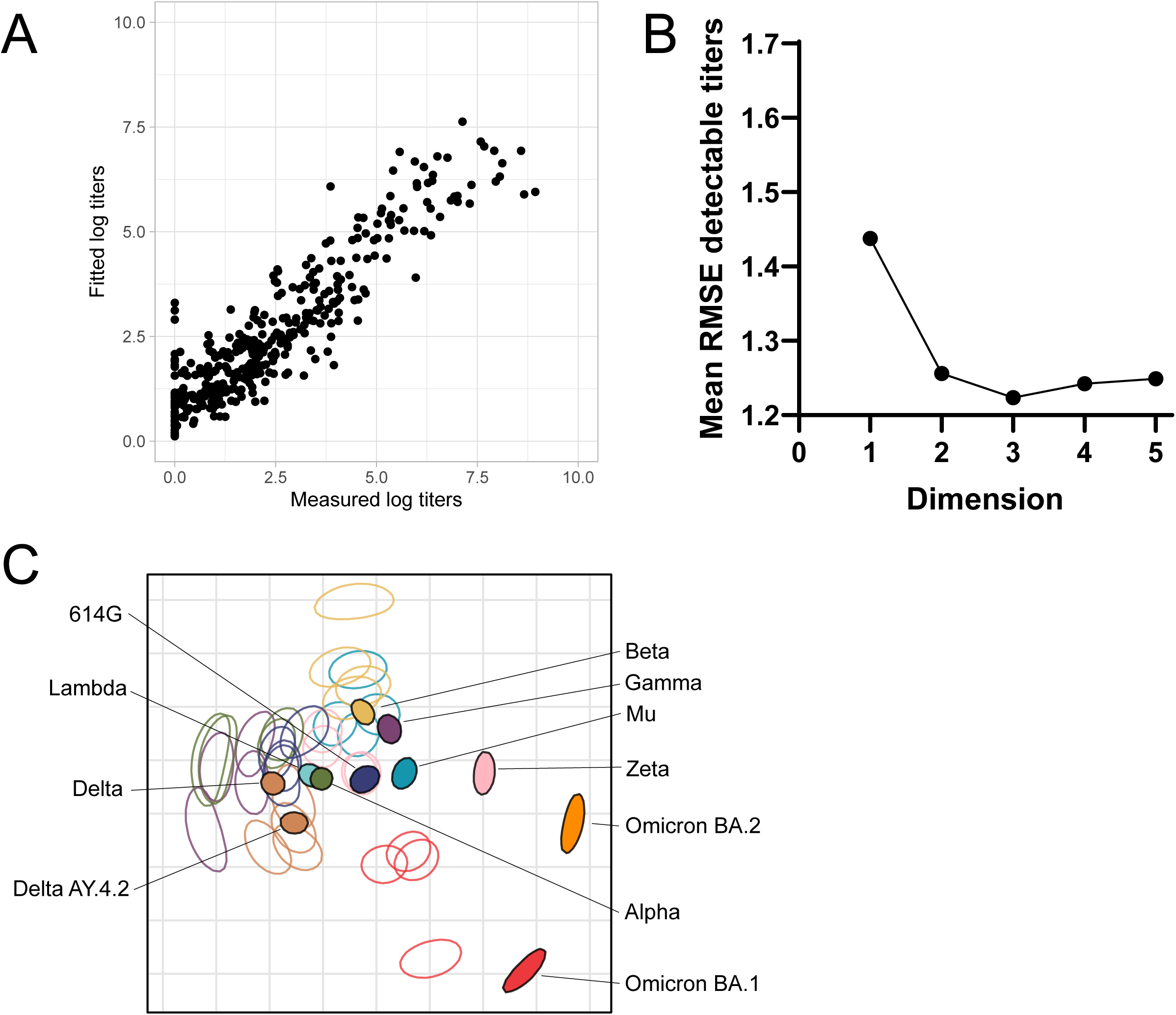
Uncertainty in antigen and antisera positions of antigenic map generated with authentic SARS-CoV-2. **a**, Scatter plot of detectable neutralizing titers on the x-axis and fitted titers from the antigenic map on the y-axis of map generated in Fig. 4i. **b**, Dimensionality test indicating the mean RMSE of detectable neutralization titers in 1 to 5 dimensions for antigenic map generated in Fig. 4i. See Extended Fig. 4 for details. **c**, Antigenic map from Fig. 4i with depicted regions (triangulation blobs) in which a particular antigen or antisera can be positioned without increasing the total stress of the map above one unit. See Fig. 3 legend for details.

